# Climate change introduces threatened killer whale populations and conservation challenges to the Arctic

**DOI:** 10.1101/2023.11.25.568606

**Authors:** Colin J. Garroway, Evelien de Greef, Kyle J. Lefort, Matt J. Thorstensen, Andrew D. Foote, Cory J. D. Matthews, Jeff W. Higdon, Caila E. Kucheravy, Stephen D. Petersen, Aqqalu Rosing-Asvid, Fernando Ugarte, Rune Dietz, Steven H. Ferguson

## Abstract

The Arctic is the fastest-warming region on the planet, and sea ice loss has opened new habitat for sub-Arctic species such as the killer whale (*Orcinus orca*). As apex predators, killer whales can cause significant ecosystem-scale changes, however, we know very little about killer whales in the Arctic. Setting conservation priorities for killer whales and their Arctic prey species requires knowledge of their evolutionary history and demography. We found that there are two highly genetically distinct, non-interbreeding populations of killer whales using the eastern Canadian Arctic—one population is newly identified as globally distinct. The effective sizes of both populations recently declined, and both are vulnerable to inbreeding and reduced adaptive potential. Furthermore, we present evidence that human-caused mortalities, particularly ongoing harvest, pose an ongoing threat to these populations. The certainty of substantial environmental change in the Arctic complicates conservation and management significantly. Killer whales bring top-down pressure to Arctic food webs, however, they also merit conservation concern. The opening of the Arctic to killer whales exemplifies the magnitude of complex decisions surrounding local peoples, wildlife conservation, and resource management as the effects of climate change are realized.

## 1. Background

The Arctic is warming four times faster than the global average[1], thus we expect to see the earliest and most significant effects of climate change in Arctic ecosystems. The increase in spatial extent and duration of the Arctic Ocean’s ice-free season is of particular concern. All credible emission scenarios predict that Arctic summers will be ice-free by the mid-20th century[2]. This will cause substantial ecosystem disruption by threatening endemic Arctic species that rely on the seasonal features of Arctic environments and by forcing northern communities to adapt to new conditions[3,4]. For example, reduced sea ice coverage increases access to the region’s resources, leading to human population growth and increasing pollution and shipping-related disturbances[5,6]. The loss of habitat for sea ice-dependent algae and phytoplankton results in negative bottom-up threats to unique sea ice ecosystems as these species underpin primary productivity at the base of the Arctic food web[3,7]. Finally, sea ice loss can result in considerable top-down pressure on Arctic food chains by providing access to predators not previously common in the region[8]. We must build our understanding of how climate change affects Arctic ecosystems to maintain what we can for people and wildlife and to hone our understanding of the complexities of ecosystem changes expected to become prevalent globally in the coming years.

Killer whales (*Orcinus orca*) are top predators with documented cascading effects on ecosystems[9–12]. The frequency of killer whale sightings in the eastern Canadian Arctic has increased considerably since the 1950s[13,14], so there is potential for them to induce substantial ecosystem-level change in this sensitive region[15]. Inuit knowledge indicates that small numbers of killer whales have long used the Arctic, but it is uncertain whether the increase in sightings is due to a growing population, more individuals from elsewhere using the Arctic, or both[15]. In any case, the growing presence of killer whales in the Arctic is likely climate-linked, as sea ice blocks access to the region for these animals[8]. With increasingly prolonged open-water seasons and growing numbers of killer whales in the region, the direct consumption of Arctic marine mammals is expected to rise[16–18]. In addition to directly affecting prey numbers, killer whales in the Arctic disrupt their prey’s habitat use and behavior, which stands to reshape marine mammal distributions[19]. Conservation efforts require information on the origin of killer whales using the Arctic, their abundance, demographic trends, population structure, and threats to their persistence to understand the consequences of global change for Arctic ecosystems.

In this study, we used whole-genome data to explore the population structure and origins of Arctic killer whales and their contemporary effective population sizes. We also conducted a comprehensive survey of published killer whale mortalities in the western North Atlantic to explore possible anthropogenic causes of recent declines. By examining population genomics and threats in Arctic killer whales, we can better understand the consequences of climate warming for Arctic ecosystems and guide future conservation and management.

## 2. Methods

### (a) Sample collection and sequencing

Killer whale samples were collected in the western North Atlantic (all samples and locations listed in table S1) through tissue biopsies from free-ranging killer whales (*n* = 20), tissue samples from harvested animals (*n* = 6), and teeth from fatally stranded killer whales (*n* = 3). We extracted DNA from skin tissue using a Qiagen DNeasy Blood and Tissue extraction kit, and DNA from tooth samples using a QIAamp DNA Investigator kit (Valencia, CA, USA). Sequencing libraries were built using sheared DNA extracts using NEBNext Ultra II DNA Library Prep Kit for Illumina (Ipswich, MA, USA) and sequenced on the Illumina HiSeq X platform (San Diego, CA, USA).

Sequencing read preparation and mapping were conducted following Foote et al.[20]. Briefly, we trimmed reads with Trimmomatic v0.35[21], then mapped the reads to a high-quality reference genome assembly (accession #GCA_000331955.1[22]) using BWA v0.7.12[23]. GATK v3.7.0[24] was used to create an interval file for suspect indels, combined with high-confidence single-nucleotide polymorphism (SNP) positions, and filtered to include only autosomal regions. We masked repeats and low-quality regions using BEDtools v2.27.1[25], then merged the aligned reads with Picard[26]. Finally, to identify genomic variants with the reference genome, we used Freebayes v1.2.0, which is a haplotype-based variant detector[27]. Variants were filtered with Vcftools v0.1.17[28] to remove indels, low quality sites (quality < 30), sites out of Hardy-Weinberg Equilibrium (p-value threshold < 0.005), and sites with missingness > 0.4. SNPs were further filtered for minor allele frequency < 0.05 and pruned for linkage disequilibrium (LD r^2^ > 0.8) to create a dataset for population structure analyses. See supplemental materials for further details on sample collection, DNA extraction, and sequencing reads.

### (b) Genomic analyses

First, we estimated kinship through R-package SNPRelate v1.30.1[29] and plink v1.9[30], and removed duplicates and one individual from each close kin pair in downstream analyses (removed individuals are marked in table S1). Next, to assess Arctic killer whale population structure, we used Principal Component Analysis (PCA) with R-package adegenet v2.1.7[31,32] and examined ancestral admixture through sparse non-negative matrix factorization (sNMF) in the R-package LEA v3.8.0[33]. We used R-package StAMPP v1.6.3[34] to calculate a fixation index (*F*_ST_) to measure genetic differentiation between the two putative populations (High Arctic and Low Arctic). Following parameters used in Foote et al.[35], we examined runs of homozygosity (ROH) across individual genomes using plink v1.9[30] then measured the frequency of ROH across the genome.

To place the Arctic killer whales within a global context of killer whale populations, we compared individuals from the two Arctic populations to genomes sampled from 25 additional sites worldwide[20]. Associations among all samples were investigated using PCA, then plotted as a covariance matrix. ABBA BABA statistics (Patterson’s D statistics) were used to test for introgression[36,37] among High and Low Arctic, and samples in the global dataset (X) at sites where X had a derived allele (i.e., an allele different to that in an ancestral outgroup).

Demographic history was assessed with several methods to examine historic and contemporary effective population sizes (*N*_e_). Using SMC++ v1.15.2[38], we identified historic changes in *N*_e_ over time and estimated when the two killer whale populations diverged. Here, we used a mutation rate of 2.34 x 10^-8^ based on Dornburg et al.[39] and a generation time of 25.7 years for this species[40]. To estimate recent changes in *N*_e_ (within the last 150 generations), we used a linkage-based method in the program GONE[41] and followed parameters used in Kardos et al.[42]. Finally, we estimated contemporary effective population sizes for both populations separately through StrataG v2.5.1[43], using a SNP dataset that was further filtered and randomly down-sampled to 25,000 SNPs.

For further detail on genomic analyses methods, please refer to the supplemental materials.

### (c) Survey of anthropogenic killer whale mortalities

To assemble a database of killer whale mortalities from anthropogenic causes in the western North Atlantic, we searched for records of these events bounded by 45°W longitude (south of Cape Farwell, Western Greenland) and 10°N latitude (to Trinidad and Tobago). Mortality types were divided into categories for commercial whaling, subsistence harvests, opportunistic harvests, fishing gear entanglement, and retaliatory.

Some uncertainty in the number of records could be due to some sources pooling catch records by month, a likely underestimated number of commercial kills, potential under-reported struck and loss rates, and discrepancies between sources. We did not include mortality from natural causes.

## 3. Results and discussion

### (a) Population structure and evolutionary origins of Arctic killer whales

We found clear, consistent evidence for two genetically distinct populations using the eastern Canadian Arctic. One population comprised individuals sampled in the eastern Canadian High Arctic and Newfoundland (hereafter referred to as the “High Arctic” population); the second included individuals sampled from the Canadian Low Arctic and Greenland (hereafter referred to as the “Low Arctic” population) (figure 1). Although the geographical range of these two populations overlaps temporally in the Arctic, they were highly genetically distinct (*F*_ST_ = 0.198; 0.198 to 0.200 95% CI). When comparing the Arctic whales with the global dataset, High Arctic killer whale genomes were distinct with this analysis as well, and Low Arctic whales were genetically similar to individuals sampled from the eastern North Atlantic (Greenland, Norway, and Iceland) (figure 2). These analyses suggest that there is very limited or no contemporary gene flow between the two groups. Notably, the High Arctic individuals were genetically distinct from all other sampled populations—they thus comprise a newly genetically identified population of killer whales. It is possible that the High Arctic population is the Arctic population Inuit communities have long seen in the region, but genetic confirmation of this would require older samples than are currently available.

**Figure 1.**
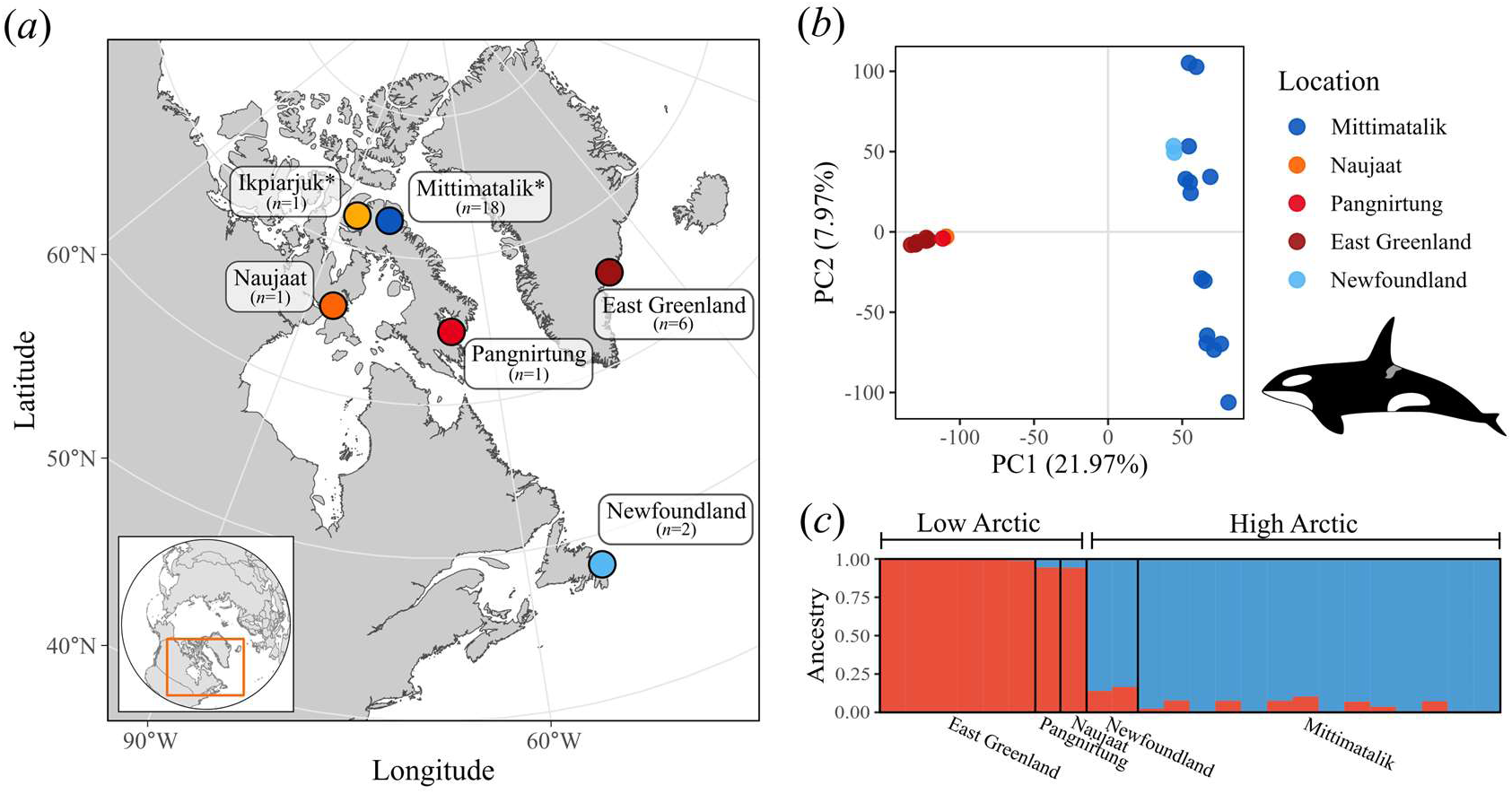
Two sympatric killer whale populations in the eastern Canadian Arctic. (*a*) Sampling locations with respective sample sizes in parentheses (*n* = 29). Locations with an asterisk (*) include individuals from close kin or duplicate pairs that were excluded in population structure analyses. (*b*) PCA containing high proportion of variance and (*c*) admixture results support evidence of two genetic populations.

**Figure 2.**
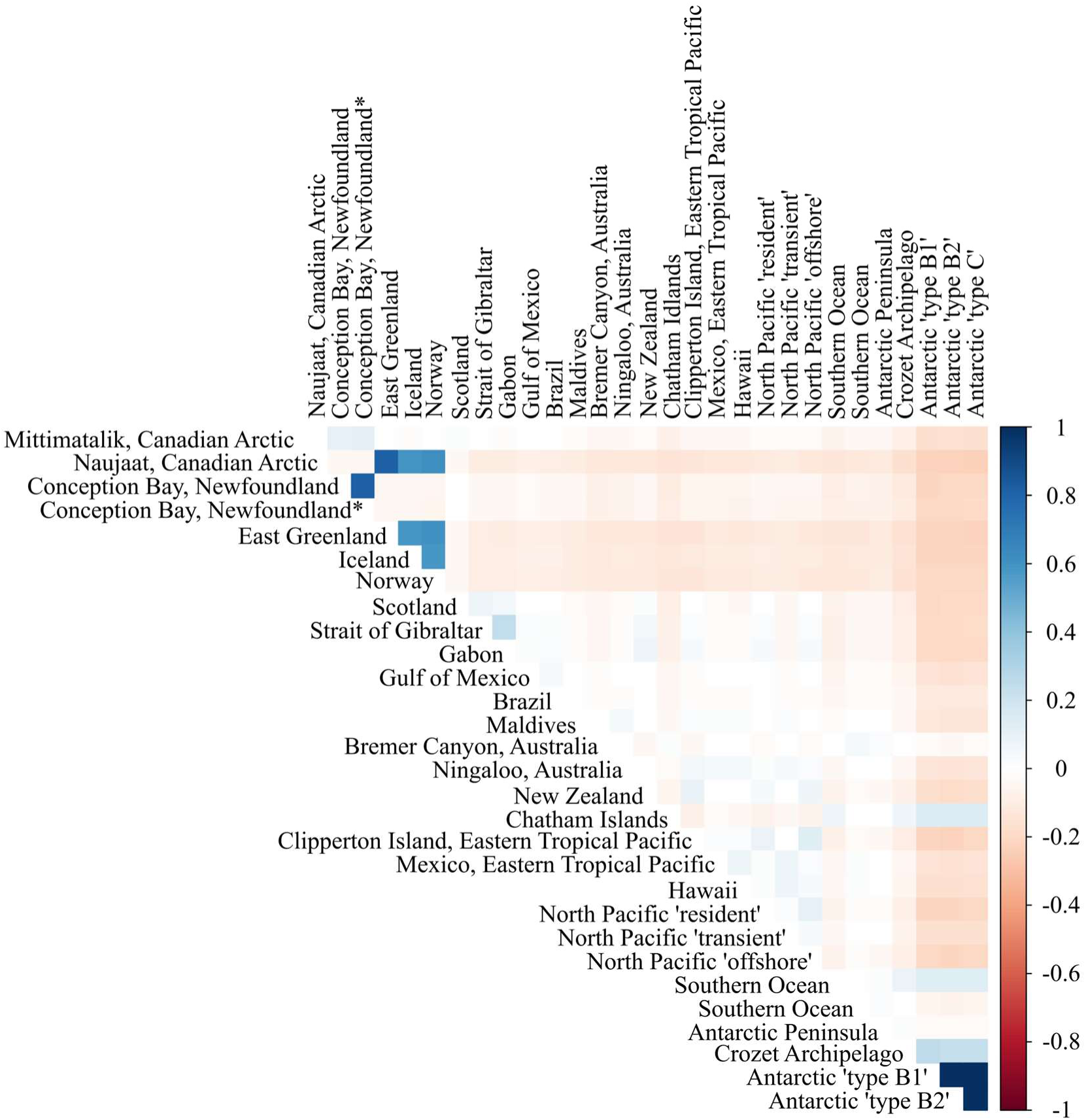
Arctic killer whales positively covaried with whales sampled in Newfoundland and eastern North Atlantic. Covariance matrix from a PCA among global killer whales with two samples from this study and samples from a previously published global dataset in Foote et al. (2019). “Mittimatalik, Canadian Arctic” represents the High Arctic population; and “Naujaat, Canadian Arctic” represents the Low Arctic population.

The evolutionary origins of the High and Low Arctic killer whales suggest their co-occurrence in eastern Canadian Arctic waters is a secondary contact between an ancestral Atlantic population and a derived sub-Arctic population. Analyses of shared ancestry among the global sample of killer whales suggest that High Arctic killer whales are derived from an ancestral Atlantic population; they harbored an excess of derived alleles shared with killer whales sampled in Brazil and Newfoundland. In contrast, Low Arctic killer whales likely derived from a population that expanded into the eastern North Atlantic from Greenland, Iceland, and Norway (table S2; figure S1). We estimated the timing of divergence between the High and Low Arctic populations to have occurred near the end of the Last Glacial Maximum approximately 9–20 kya ago (figure S2). The lack of strong genetic similarity and shared ancestry with killer whales outside the Atlantic Ocean suggest that these two populations evolved within the Atlantic rather than via colonization from a different region.

We know relatively little about the ecology of the killer whales that use the Arctic[15]. However, ecological divergence and specialization underlie genetic differentiation between killer whale populations elsewhere. For example, killer whales in the northeastern Pacific Ocean and the Southern Ocean have recognizable ecotypes based on diet, social behavior, morphology, and genetics[44–49]. The level of differentiation between the High and Low Arctic killer whales is comparable to the difference between ecotypes from within the Antarctic and Pacific oceans[50]. However, it is much less clear whether North Atlantic populations can be ecologically categorized as discretely[51,52]. Fatty acid signatures from killer whales and their prey point to gradients in diet across the region—killer whales in the western North Atlantic, including Canadian waters, primarily consumed other whale species (e.g., [53]), and individuals from Greenland prey mainly on seals and fish such as herring (*Clupea harengus)* and mackerel (*Scomber scombrus*)[54]. Direct observations of killer whales feeding in the Canadian Arctic support these findings, with beluga (*Delphinapterus leucas*) and narwhal (*Monodon monocero*s) observed as prey most often, followed by bowhead whales (*Balaena mysticetus*), ringed (*Pusa hispida*), harp (*Pagophilus groenlandicus*), bearded (*Erignathus barbatus*), and hooded (*Cystophora cristata*) seals[13,55]. In Greenland, Inuit hunters report seals as the main killer whale prey, followed by fish and minke whales and narwhals in the northernmost part of West Greenland[56–58]. Therefore, we have limited data to support ecological discreteness, but strong evidence for genetic differentiation in killer whale populations of the eastern Canadian Arctic.

### (b) Population demography and threats to Arctic killer whales

The most important quantities for understanding population biology, and for conservation and management decision-making, are the number of individuals in a population, the effective population size, and whether these values are trending up or down. The number of individuals in a population governs population ecology, and the effective population size shapes several evolutionary processes[59]. The effective population size is an estimate of the strength of genetic drift a population experiences. The smaller the effective population size, the faster a population loses genetic diversity. This is important because genetic diversity contributes to population mean fitness and the capacity to adapt to current and future environmental change[60]. Additionally, the efficiency of natural selection is inversely proportional to the strength of drift, meaning small populations will have difficulties adapting to environmental change. In practice, the effective population size will be much smaller than, and not well correlated with, the census population size due to variation in reproductive success and output across individuals and the population’s demographic history. For conservation and management, it is important to note that population sizes increase much faster than genetic diversity due to the slow rate at which mutations accrue. Thus, growing and even relatively large populations can be at risk, evolutionarily speaking, if their effective size is low. Below we explore threats to Arctic killer whale populations related to trends in effective population size and the numbers of individuals in the populations.

The genetic diversity and demographic histories of the Canadian Arctic killer whale populations strongly suggest that they warrant conservation concern. We found the effective population sizes of the High and Low Arctic populations are 20 (*N*_e_ = 19.67; 19.65 to 19.69 95% CI) and 14 (*N*_e_ = 13.92; 13.89 to 13.94 95% CI), respectively. These very small effective population sizes mean that these populations will have difficulty adaptively responding to future environmental change. Recently, the United Nations Convention on Biodiversity adopted the Kunming-Montreal global biodiversity framework to guide biodiversity conservation interventions. This agreement prioritized two indicators of population genetic risk. First, the agreement considers effective population sizes < 500 to be at high genetic risk due to loss of adaptive capacity [headline indicator A.5 for Goal A and Target 4CBD, 2022[61]]. The effective sizes identified here are approximately 14x to 20x below United Nations guidelines for limiting genetic risk. While the relationships between effective population size and genetic risk are context-dependent, the low effective population sizes in the High and Low Arctic are concerning for their capacities to maintain genetic diversity and adaptive resilience. The second headline indicator for evolutionary risk is the loss of genetically distinct populations. The genetically distinct Arctic populations we identify merit conservation and management concern based on both criteria.

The two Arctic populations arrived at their similarly low effective population sizes differently and in ways that will affect conservation efforts and likely their outcomes. The populations drifted genetically following their initial divergence (figure S2) and the effective population sizes of both populations were stable for most of the last 4000 years (figure 3a). However, both populations have experienced notable recent declines (figure 3a). The High Arctic population decline was much more pronounced than in the Low Arctic population, which had a much smaller effective population size across those 4000 years. Although both populations’ contemporary effective sizes are similar, the Low Arctic population is much more inbred than the High Arctic population, as shown by the greater proportion of runs of homozygosity (figure 3b). The difference in inbreeding between the High Arctic and Low Arctic is likely due to the recency and magnitude of the decline in effective population size in the High Arctic whales, as genetic drift and inbreeding accumulate for many generations after the initial population declines. Given the similar contemporary effective population sizes, we should expect ongoing inbreeding in the Low Arctic population with the levels of genetic diversity in both populations eventually converging.

**Figure 3.**
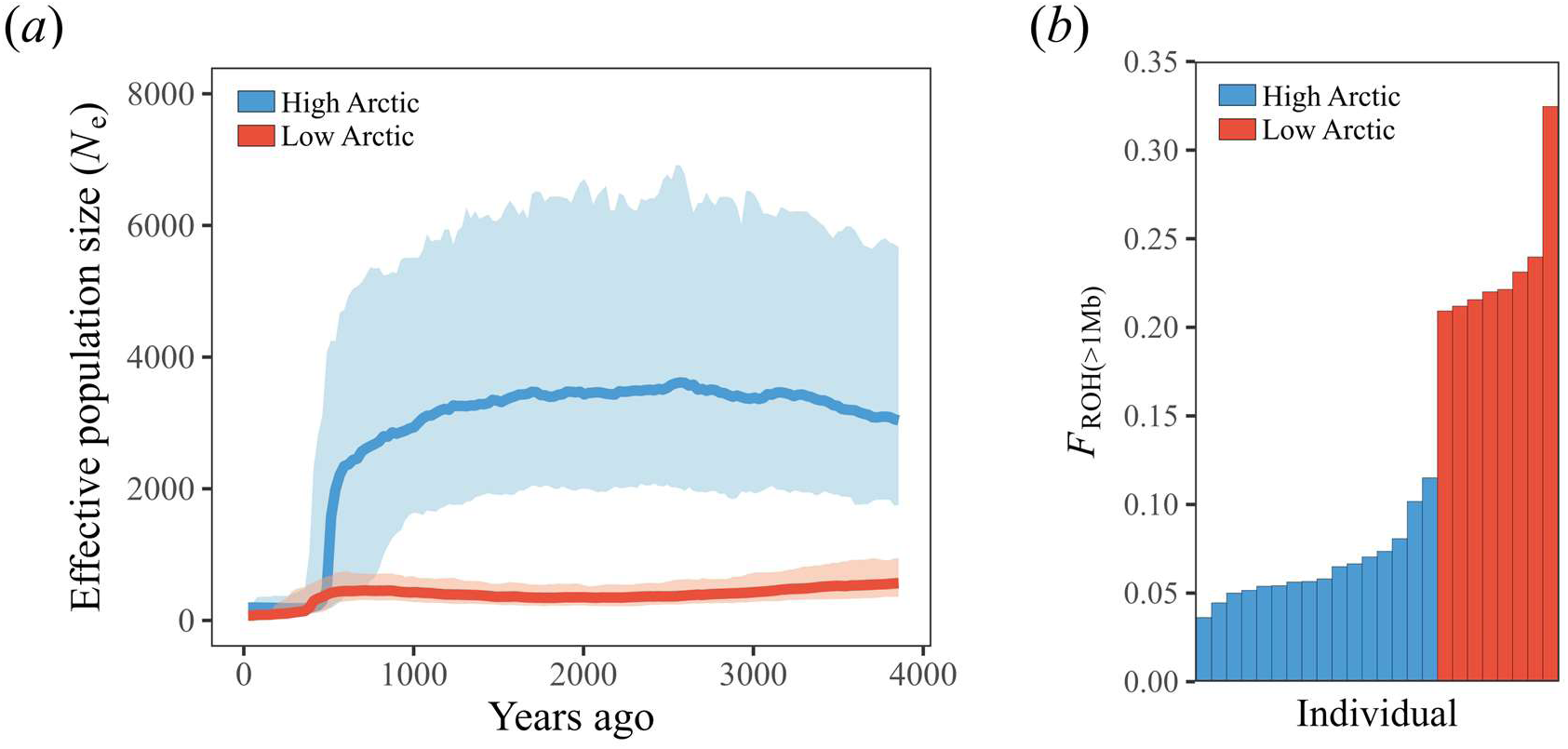
Recent population declines and levels of inbreeding. (*b*) Arctic killer whale effective population sizes across over the past 150 generations (generation time of 25.7 years[40]) (see figure S3 for the model without minor allele frequency filter). Bolded lines represent the median estimates and shaded regions are 95% confidence intervals. (*b*) Proportions of runs of homozygosity (ROH) using a minimum ROH length of 1 Mb for each killer whale individual from the eastern Canadian Arctic (see figure S4 with a minimum ROH length of 1.5 Mb).

The very low contemporary effective population sizes suggest that Arctic killer whales are vulnerable to inbreeding depression. Inbreeding depression causes reduced survival and reproduction in small populations due to increased homozygosity of partially recessive deleterious alleles. The well-documented negative effects of inbreeding in Southern Resident killer whales are instructive for the Arctic populations[42]. The Southern Resident and Low Arctic populations have very comparable recent demographic histories and patterns of runs of homozygosity, and all three populations have comparable contemporary effective population sizes (Southern Resident *N*_e_ = 27, Kardos et al.[42]; High Arctic *N*_e_ = 20; Low Arctic *N*_e_ = 14). Using long-term individual based monitoring data in the Southern Resident population, Kardos et al.[42] found convincing evidence that inbreeding depression limits population growth in Southern Resident killer whales, and predicted further population declines in the system due to inbreeding. Given historical and contemporary demographic similarities with the Low Arctic population, those findings bear on the current and future effects of inbreeding for the Arctic whales—inbreeding depression will likely limit population recovery. While both Arctic populations may be threatened by inbreeding, the smaller runs of homozygosity in the High Arctic population mean that conservation action focused on extrinsic threats and maintaining population size could mitigate the risk of inbreeding depression in this population.

Census population size estimates help put effective population sizes into context regarding human-caused mortalities. The number of killer whales in the northern Baffin Island region is estimated to be between 136 to 190 individuals, based on photographic capture recapture[18]. In our survey of published killer whale mortalities in the western North Atlantic (table S3, figure S5), we found that commercial whaling and subsistence harvest were the leading documented sources of mortality. The earliest whaling record in the region was from Nuuk, West Greenland, in 1756[56,62]. Harvest has continued since then, with Greenland introducing a bounty on killer whales from 1960-1975[56]. This harvest continues to this day with a distinct increase in southeast Greenland starting in 2009, following a climate-related ecosystem shift caused by the collapse of drifting sea ice during summer and a consequent increase in the number of killer whales in the area[58,63]. Of the 500 mortalities we document, all but two (fishing gear entanglement) were intentional kills. This literature survey is very likely an underestimate of human-caused mortality. Kills in Canadian waters are under-reported but second and third-hand accounts suggest they occur[64]. Killer whales are often difficult to retrieve as they readily sink after being killed due to having relatively little blubber, decreased buoyancy in cold waters[65], and because they are or were not killed for consumption—killer whales are sometimes seen as threats to people and valued wildlife[58]. These deaths often go unreported but could be substantial in number. For example, during three hunts in Greenland, interviewed hunters reported landing 4 whales with an additional 9 whales killed, unretrieved, and unreported in harvest data[58].

Movement within a population is important to consider when looking into anthropogenic mortalities, since hunting pressure in one location may affect the population inhabiting multiple areas. Our genomic results suggest the High Arctic population range between Mittimatalik and Newfoundland and photographic recaptures support this. Two individual killer whales (DI02 and DI03) first identified near Disko Island, Greenland in 2011[66] were re-sighted near Mittimatalik, Nunavut in 2019 (ECA072 and ECA071, respectively)[67]. This evidence shows that killer whales are moving between eastern Canadian Arctic and west Greenland waters, indicating that killer whales in the High Arctic population are subject to harvest pressure in Greenland. It is likely that harvest has contributed to the recent population declines we document and is a threat to population recovery.

## 4. Conservation implications

The task of conservation given climate change presents a classic ‘wicked problem’ that will continue to play out globally. Accumulated greenhouse gas emissions and future emissions targets ensure that the planet’s ecosystems will be highly altered regardless of future mitigations. With conservation of the present state in many regions ranging from impractical to impossible we are forced to conserve what we can and manage wildlife for an uncertain future. The case of killer whales in the Arctic exemplifies the magnitude of complex decisions related to people and wildlife conservationists and managers will face as the effects of climate change are realized throughout the planet. The current small, genetically homogeneous, and potentially ecologically distinct killer whale populations in the eastern Canadian Arctic are susceptible to inbreeding and harvest, as well as a high exposure to contaminants[68]. Conservation and management issues are made more complex by the lack of foundational knowledge in these systems, exemplified by the newly identified High Arctic population. At the same time, the increasing use of the Arctic and consumption of Arctic marine mammals by killer whales could also cause significant ecosystem-scale change concurrent with other threats through trophic cascades (e.g., [9–11,69]; although see [70,71]). Arctic marine mammals use sea ice to reduce the risk of killer whale predation. With the loss of ice cover, killer whale predation could lead to severe consequences for their prey populations. The marine mammals that the killer whales hunt while in northern waters are culturally and economically important to indigenous communities, so these species also merit conservation and management concern in light of killer whale populations moving into the Arctic. Effective conservation and management of killer whale populations and their ecosystems will require a holistic approach that considers their genetic background, and complex interactions among killer whale populations, changing prey interactions, and other human-induced threats in the context of ongoing climate change. This will require collaborations among scientists, policymakers, and stakeholders across national and international borders. Finally, it will require a commitment to address the root causes of threats to killer whale populations, including climate change and human activities such as past and current whaling.

## Acknowledgments

We thank the many Canadian Arctic communities and local Hunters and Trappers Organizations involved in sample collection. Two of the teeth used in this study were provided by the Manitoba Museum (Winnipeg, MB) and W. Ledwell (Portugal Cove, NL). Canadian Arctic biopsy collections were approved by the DFO Freshwater Institute Animal Care Committee AUP# FWI-ACC-2013-022 (2013) and AUP# FWI-ACC-2018-008 (2018), and permitted under DFO License to Fish for Scientific Purposes #S-13/14-1024-NU (2013) and #S18/19-1029-NU (2018). This research was enabled in part by support provided by the Prairies DRI and the Digital Research Alliance of Canada (alliancecan.ca). We also thank Denise Tenkula for DNA extraction work, Zoltan Nemeth for identifying killer whale matches, and Hal Whitehead and Levi Newediuk for feedback on the manuscript.

