## Supplementary Material for "Climate change introduces threatened killer whale populations and conservation challenges to the Arctic"

**Methods**

Sample collection

We collected killer whale samples throughout the eastern Canadian Arctic (all samples and locations listed in table S1). Tissue biopsies, comprising skin and blubber, were collected from free-ranging killer whales using a 150 lbs draw-weight Excalibur Crossbow and crossbow bolt equipped with a sterile tubular stainless steel biopsy tip (*n* = 20). Additional tissue samples were collected from animals harvested in East Greenland (*n* = 6). Teeth were collected opportunistically from fatally stranded killer whales (*n* = 3).

DNA extraction

We extracted DNA from skin tissue (10-15 mg per sample) using a Qiagen DNeasy Blood and Tissue extraction kit (Valencia, CA, USA) following the Purification of Total DNA from Animal Tissues (Spin-Column Protocol) protocol. For tooth samples, pulp powder (or dentine powder from the surface layer of the root when pulp was not available) was first collected with a hand drill and 3 mm ball-shaped diamond-coated drill bits surface-cleaned with ELIMINase Decontaminant. We extracted DNA from the powdered tooth (~100 mg) using a QIAamp DNA Investigator kit (Valencia, CA, USA) following the Isolation of Total DNA from Bones and Teeth protocol. Genomic DNA was sheared to ~150 bp using a Covaris LE220 (Woburn, MA, USA). Sequencing libraries were built using sheared DNA extracts using NEBNext Ultra II DNA Library

Prep Kit for Illumina (Ipswich, MA, USA) and sequenced on the Illumina HiSeq X platform (San Diego, CA, USA).

Whole-genome sequencing

Sequencing read trimming, mapping, filtering, and repeat-masking were conducted following Foote et al.[1]. We processed reads with Trimmomatic v0.35[2] to remove adapter sequences, trailing low-quality regions (Q < 20), and N bases, and to drop reads < 31 bases long. We then mapped the reads to a high-quality reference genome assembly (accession #GCA_000331955.1[3]). Processed reads were aligned to the mitochondrial genome using BWA v0.7.12[4]. Unmapped reads were extracted using SAMtools v1.5[5] and aligned to the nuclear genome using BWA[4]. GATK v3.7.0[6] was used to create an interval file for suspect indels, combined with high-confidence single-nucleotide polymorphism (SNP) positions, and filtered to include only autosomal regions. Repeats and low-quality regions were masked using BEDtools v2.27.1[7]. We merged the aligned reads with Picard[8] and then identified genomic variants with the reference genome using Freebayes v1.2.0 which is a haplotype-based variant detector[9]. Variants were filtered with Vcftools v0.1.17[10] to remove indels, low quality sites (quality < 30), sites out of Hardy-Weinberg Equilibrium (p-value threshold < 0.005), and sites with missingness > 0.4. This resulted in 1,351,642 SNPs. SNPs were further filtered for minor allele frequency < 0.05 and pruned for linkage disequilibrium (LD $r^2 > 0.8$) to create a dataset for kinship analyses and population structure, resulting in 421,668 SNPs.

Population structure

Principal Component Analysis (PCA) was used to explore associations among samples, completed in the R-package adegenet v2.1.7[11,12]. Pairwise kinship coefficients (Φ), the probability that randomly sampled homologous alleles from two individuals were identical-by-descent (i.e., copies of the same ancestral gene), among all pairs of individuals were calculated using the Maximum Likelihood Estimation Method in the R-package SNPRelate v1.30.1[13]. Based on initial PCA results showing two genetic clusters, we estimated kinship coefficients on each cluster (High Arctic and Low Arctic) separately. We removed duplicates and one individual from each close kin pair in downstream analyses (removed individuals are marked in table S1).

We examined ancestral admixture through sparse non-negative matrix factorization (sNMF) in the R-package LEA v3.8.0[14], running models with 1 – 5 for the K value (representing the number of ancestral sources) with 10 repetitions per K value. To estimate genomic differentiation between the two putative populations, we calculated a fixation index ($F_{ST}$) with R-package StAMPP v1.6.3 through 100 bootstraps[15]. We examined runs of homozygosity (ROH) across individual genomes for each population using plink v1.9[16]. Following Foote et al.[17], we used these parameters: minimum segment length of 300 kb, minimum number of 50 SNPs, density of at least one SNP per 50 kb, gap of 1000 kb, up to 3 heterozygote sites per 300 kb windows, and allowing up to 10 missing calls in a window. We then measured the frequency of ROH across the genome with a minimum ROH length of 1 Mb ($F_{ROH>1Mb}$) and 1.5 Mb ($F_{ROH>1.5Mb}$).

Global population structure

One sample from each of the putative populations identified in this study (ARPI_2013_4001 from Mittimatalik representing the High Arctic; ARRB_2009_1291 from Naujaat representing the Low Arctic) was included in an analysis with samples representing the species' global range. We randomly sampled an allele at each SNP from the genomes to account for variation in coverage. Associations among all samples were investigated using PCA, then plotted as a covariance matrix. ABBA BABA statistics (Patterson's D statistics) were used to test for introgression[18,19] among High and Low Arctic, and samples in the global dataset (X) at sites where X had a derived allele (i.e., an allele different to that in an ancestral outgroup). For the ancestral outgroup, we used the consensus genome from the common bottlenose dolphin (*Tursiops truncatus*) and the long-finned pilot whale (*Globicephala melas*) as the representative of the ancestral allele at each site[1]. We compared the number of sites where High Arctic (and not Low Arctic) shared a derived allele with X, and the number of sites where Low Arctic (and not High

Arctic) shared a derived allele with X. This test was repeated for each sample in the global dataset (see [1] for details).

<u>Demographic history</u>

SMC++ v1.15.2 was used to identify when western North Atlantic killer whale groups split and approximate effective population size (*N*e) over time[20]. Biallelic SNPs above with genotype and site quality ≥30 were used for this analysis, but because runs of homozygosity affect demographic reconstruction with SMC++, indels and unused SNPs were also provided as masked sites—this file was created with bedops v2.4.39[21]. For the two populations, composite likelihoods were created by using a distinguished individual from each lineage within each population. For group one (High Arctic), individuals 48335KWEG2012, ARRB_2009_1291, and Pang2013 were used as distinguished pairs. For group two (Low Arctic), individuals ARPI_2013_4001, B045, D118_2, and OO_01 were distinguished pairs. Input files were created with for each distinguished pair, per contig. The estimate was used in 100 replicates for each population with a mutation rate of $2.34 \times 10^{-8}$ based on Dornburg et al.[22]. In addition, a regularization penalty of 4.0, a non-segregating site cutoff of 100,000, thinning of 2,000 and timepoints of 10 to 10,000 generations were used. The 100 replicate model outputs per population were combined into 100 combined models with both populations using SMC++'s split function. All 100 replicates were plotted together, using a generation time of 25.7 years for this species[23]. SMC++ v1.15.2 was used to identify when western North Atlantic killer whale groups split and approximate *N*e over time[20]. Biallelic SNPs above with genotype and site quality ≥30 were used for this analysis, but because runs of homozygosity affect demographic reconstruction with SMC++, indels and unused SNPs were also provided as masked sites—this file was created with bedops v2.4.39[21]. For the two populations, composite likelihoods were created by using a distinguished individual from each lineage within each population. For group one (High Arctic), individuals 48335KWEG2012, ARRB_2009_1291, and Pang2013 were used as distinguished pairs. For group two (Low Arctic), individuals ARPI_2013_4001, B045, D118_2, and OO_01 were distinguished pairs. Input files were created with for each distinguished pair, per contig. The estimate was used in 100 replicates for each population with a mutation rate of 2.34 x $10^{-8}$ based on Dornburg et al.[22]. In addition, a regularization penalty of 4.0, a non-segregating site cutoff of 100,000, thinning of 2,000 and timepoints of 10 to 10,000 generations were used. The 100 replicate model outputs per population were combined into 100 combined models with both populations using SMC++'s split function. All 100 replicates were plotted together, using a generation time of 25.7 years for this species[23].

To estimate recent demographic history (within the last 150 generations), we used a linkage-based method in the program GONE[24]. Here, we ran models with and without the minor allele frequency threshold, and each population separately. Following Kardos et al.[25], we used input parameters of 1000 generations for linkage data obtains in bins, 1000 bins, maximum numbers of 10,000 SNPs per scaffold, an hc value (recombination rate) of 0.02, and 500 repetitions.

We estimated contemporary effective population sizes for both populations separately through StrataG[26]. Using Vcftools[10] and plink v1.9[16], SNPs were further filtered for no missing data, minor allele frequency > 0.05, and randomly down-sampled to 25,000 SNPs. We used three separate subsets of 25,000 SNPs to reduce bias from random chance. We estimated contemporary effective population sizes for both populations separately through StrataG v2.5.1[26]. Using Vcftools v0.1.17[10] and plink v1.9[16], SNPs were further filtered for no missing data, minor allele frequency > 0.05, and randomly down-sampled to 25,000 SNPs. We used three separate subsets of 25,000 SNPs to reduce bias from random chance.

### Figures

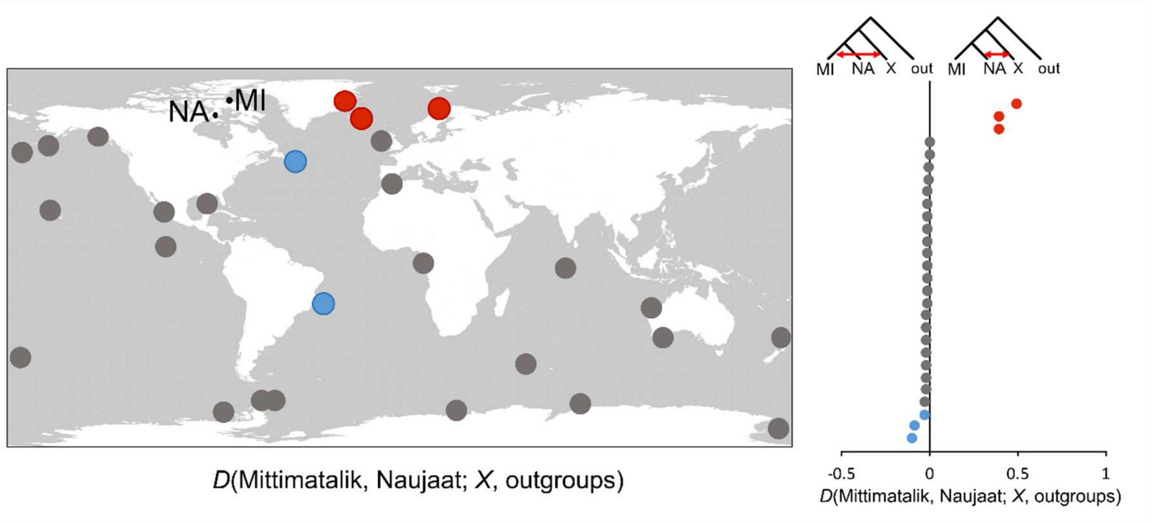


**Figure S1.** Whole-genome sequenced killer whales included in the global ABBA BABA analysis, with site locations from Foote et al. (*20*) and Naujaat (NA) and Mittimatalik (MI) in the map (left). Points are colour-coded based on the values of *D*(Mittimatalik, Naujaat; X, outgroup) (right). Blue points represent significant Z-scores < -3, which indicates an excess of derived alleles shared with Mittimatalik, and red points represent significant Z-scores > 3, which indicates an excess of derived alleles shared with Naujaat (results also shown in Table S2).

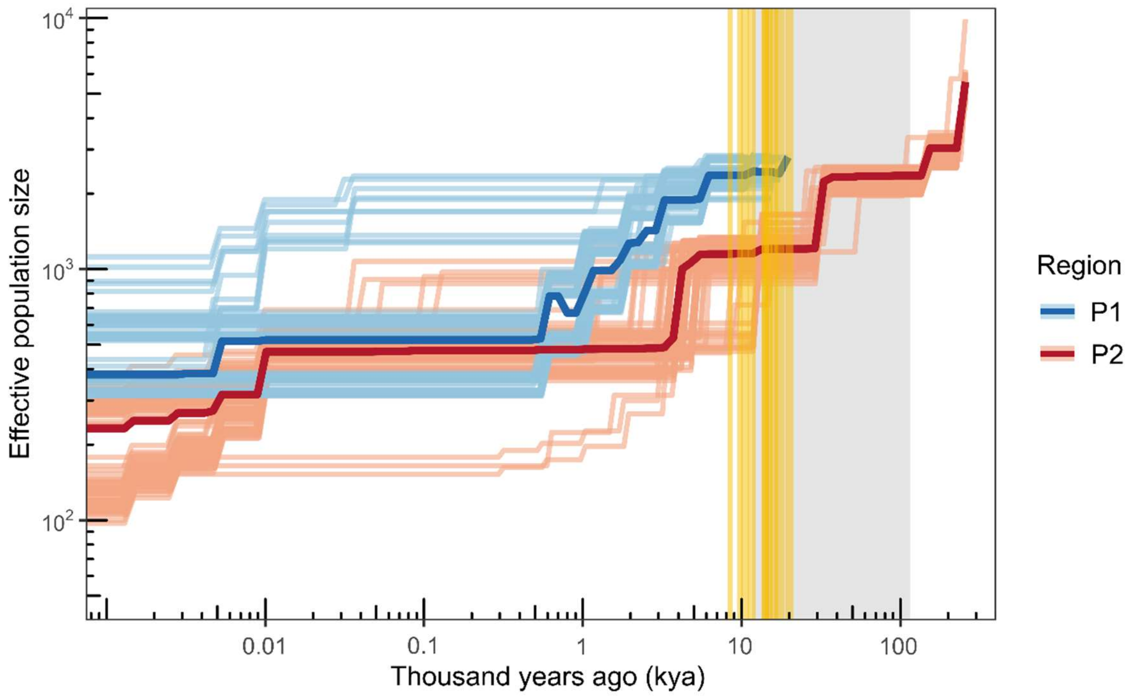


**Figure S2.** Demographic history of eastern Canadian Arctic killer whales, showing an overall decline in effective population size over past time and a population divergence (yellow vertical lines estimated at 9-20 kya) near the end of the last glacial period (11.7-115 kya; shaded in gray). P1 (*n* = 16; High Arctic) is shown in blue, and P2 (*n* = 8; Low Arctic) is shown in red. The darker line in each group represents the median estimate among 100 iterations.

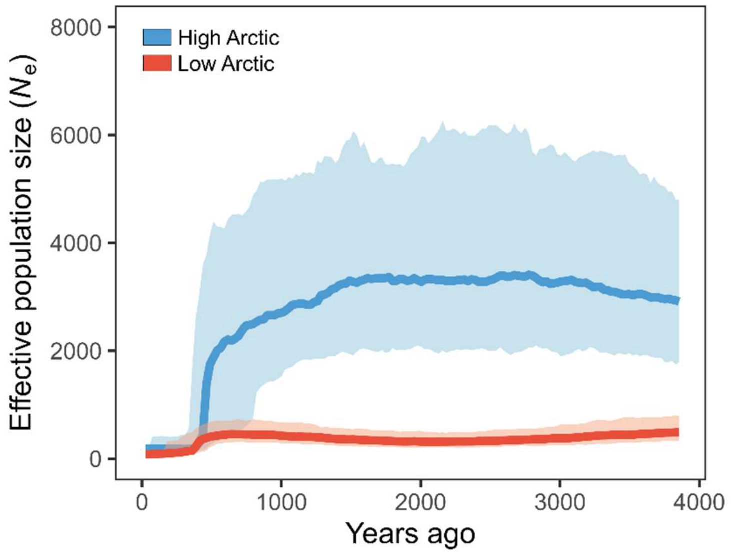


**Figure S3.** Arctic killer whale effective population sizes across over the past 150 generations (using generation time of 25.7 years) excluding a minor allele frequency filter.

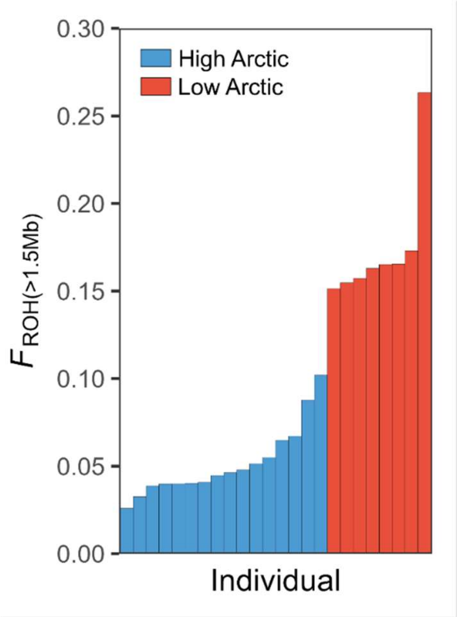


**Figure S4.** Proportions of runs of homozygosity (ROH) using a minimum ROH length of 1.5 Mb for each killer whale individual (*n* = 24).

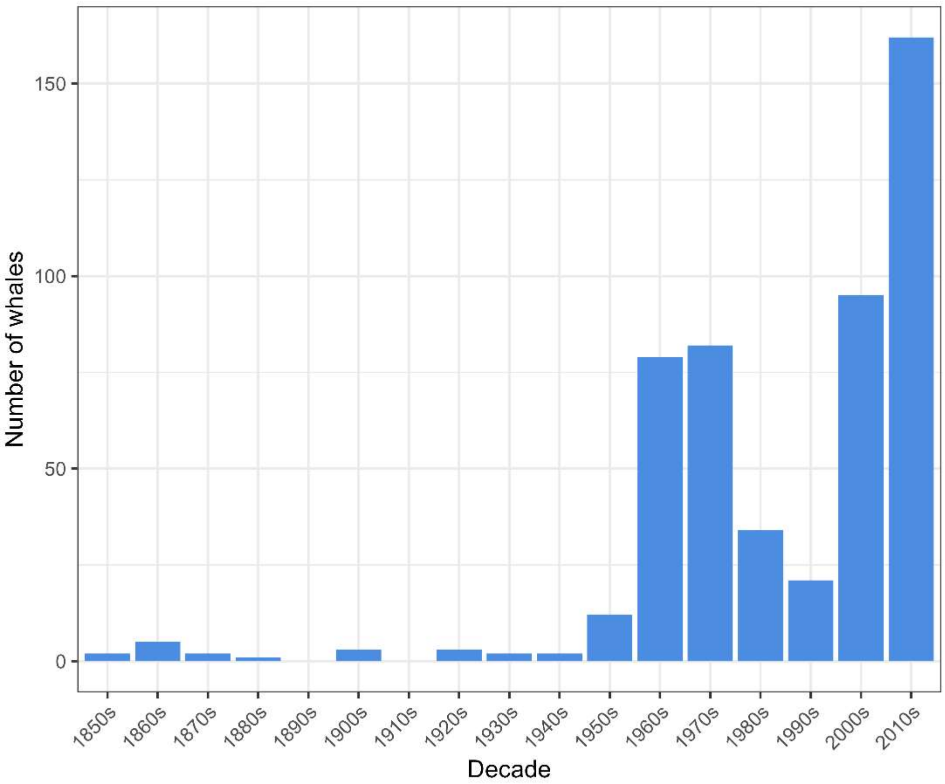


**Figure S5.** Decadal summary of recorded number of killer whale deaths caused by anthropogenic causes in the western North Atlantic.

**Table S1.** Sample information including ID, location, collection year, type of killer whale samples, and mean coverage of genomic data. Duplicates and close kin pairs that were removed from population structure and demographic history analysis are marked with an X and the ID of the related individual. Samples used in the global analysis are marked in the last column.

| Sample ID | Location | Year | Sample type | Mean coverage | Global analysis | Samples removed |
|---|---|---|---|---|---|---|
| MM406_2 | Ikpiarjuk | 1948 | tooth | 13.9 | | X – pair with ARRB_2009_1291 |
| D118_2 | Newfoundland | 1971 | tooth | 15.0 | | |
| B045 | Newfoundland | 2002 | tissue | 10.3 | | |
| ARRB_2009_1291 | Naujaat | 2009 | tooth | 16.2 | X | |
| 48335KWEG2012 | East Greenland | 2012 | tissue | 14.3 | | |
| 48336KWEG2012 | East Greenland | 2012 | tissue | 16.2 | | |
| 48337KWEG2012 | East Greenland | 2012 | tissue | 12.5 | | |
| 48338KWEG2012 | East Greenland | 2012 | tissue | 13.9 | | |
| 48339KWEG2012 | East Greenland | 2012 | tissue | 15.7 | | |
| 48340KWEG2012 | East Greenland | 2012 | tissue | 15.3 | | |
| ARPI_2013_4001 | Mittimatalik | 2013 | tissue | 16.6 | X | |
| ARPI_2013_4002 | Mittimatalik | 2013 | tissue | 14.7 | | |
| ARPI_2013_4003 | Mittimatalik | 2013 | tissue | 14.1 | | |
| ARPI_2013_4004 | Mittimatalik | 2013 | tissue | 16.4 | | X – pair with ARPI_2013_4004 |
| ARPI_2013_4005 | Mittimatalik | 2013 | tissue | 14.5 | | |
| ARPI_2013_4006 | Mittimatalik | 2013 | tissue | 15.5 | | |
| ARPI_2013_4007 | Mittimatalik | 2013 | tissue | 15.3 | | |
| Pang2013 | Pangnirtung | 2013 | tissue | 15.3 | | |
| OO_01 | Mittimatalik | 2018 | tissue | 10.0 | | |
| OO_02 | Mittimatalik | 2018 | tissue | 12.2 | | X – pair with OO_17 |
| OO_07 | Mittimatalik | 2018 | tissue | 10.7 | | |
| OO_10 | Mittimatalik | 2018 | tissue | 9.0 | | X – pair with OO_14 |
| OO_11 | Mittimatalik | 2018 | tissue | 11.6 | | |
| OO_13 | Mittimatalik | 2018 | tissue | 9.8 | | |
| OO_14 | Mittimatalik | 2018 | tissue | 11.7 | | |
| OO_15 | Mittimatalik | 2018 | tissue | 11.1 | | X – pair with OO_16 |
| OO_16 | Mittimatalik | 2018 | tissue | 9.3 | | |
| OO_17 | Mittimatalik | 2018 | tissue | 12.8 | | |
| OO_18 | Mittimatalik | 2018 | tissue | 13.0 | | |

**Table S2.** nABBA, nBABA, jackknife-estimated D statistics, standard error, and Z scores among Low Arctic (ARRB-2009-1291) and High Arctic (ARPI-2013-4001) killer whales and samples in the global dataset (see Foote et al. 2019). Bolded samples share a significant excess of derived alleles with one of the two Canadian Arctic populations.

| Sample | nABBA | nBABA | D Statistic | SE | Z |
|---|---|---|---|---|---|
| **Greenland** | **184152** | **61715** | **0.498** | **0.0062** | **80.6499** |
| **Iceland** | **102320** | **43699** | **0.4015** | **0.0080** | **50.3432** |
| **Norway** | **102071** | **43430** | **0.403** | **0.0082** | **49.2925** |
| Scotland | 63065 | 62784 | 0.0022 | 0.0084 | 0.2672 |
| ETP-Mexico | 59801 | 59605 | 0.0016 | 0.0067 | 0.2457 |
| Antarctic Peninsula | 59642 | 59727 | -0.0007 | 0.0066 | -0.1072 |
| Chatham Islands | 57703 | 58556 | -0.0073 | 0.0069 | -1.0641 |
| Hawaii | 59858 | 60808 | -0.0079 | 0.007 | -1.1278 |
| Transient | 59781 | 60944 | -0.0096 | 0.0068 | -1.4219 |
| New Zealand | 59153 | 60515 | -0.0114 | 0.0075 | -1.5083 |
| Bremer Canyon | 58516 | 59837 | -0.0112 | 0.0066 | -1.6799 |
| Offshore | 58454 | 60056 | -0.0135 | 0.0075 | -1.8009 |
| type B1 | 52619 | 54169 | -0.0145 | 0.0078 | -1.8717 |
| type C | 53313 | 54835 | -0.0141 | 0.0074 | -1.8916 |
| Ningaloo | 59251 | 60830 | -0.0131 | 0.0066 | -1.9931 |
| Madlives | 57255 | 58904 | -0.0142 | 0.0069 | -2.0568 |
| Gabon | 60971 | 62800 | -0.0148 | 0.007 | -2.0996 |
| resident | 58175 | 60084 | -0.0161 | 0.0074 | -2.1791 |
| Southern Ocean | 57268 | 59030 | -0.0152 | 0.0067 | -2.2725 |
| ETP-Clipperton Island | 61032 | 63520 | -0.02 | 0.0081 | -2.4526 |
| Gulf of Mexico | 61491 | 63791 | -0.0184 | 0.0074 | -2.4918 |
| type B2 | 53920 | 56051 | -0.0194 | 0.0075 | -2.5978 |
| Southern Ocean | 58167 | 60418 | -0.019 | 0.0067 | -2.8166 |
| Gibraltar | 60491 | 63252 | -0.0223 | 0.0078 | -2.85277 |
| Crozer | 56319 | 58670 | -0.0204 | 0.0061 | -3.3747 |
| **Brazil** | **60320** | **63118** | **-0.0227** | **0.0067** | **-3.3889** |
| **Newfoundland** | **58790** | **70732** | **-0.0922** | **0.0084** | **-10.9827** |
| **D118_2** | **92761** | **108334** | **-0.0774** | **0.0057** | **-13.5733** |

**Table S3.** Records of killer whale anthropogenic mortality in the western North Atlantic, including year, location, number of killer whales harvested, harvest type, and source(s) for each record.

| Year | Location | Number of whales | Harvest type | Source(s) |
|---|---|---|---|---|
| 1756 | Nuuk, West Greenland | Some | Subsistence | Winge 1902; Heide-Jørgensen 1988 |
| 1840 | Disko Bay, West Greenland | Pod/Several | Subsistence | Winge 1902; Heide-Jørgensen 1988 |
| 1857 | Cumberland Sound, Nunavut | 2 | Commercial | Ross 1997: 78 |
| 1864 | North Atlantic | 1 | Commercial | D. B. Stewart, unpubl. data (logbook read by R.R. Reeves) |
| 1860s (?) | Belle Amours, Strait of Belle Isle, Quebec | 1 | Commercial | Weiz and Packard 1866, p. 272 in Mitchell and Reeves 1988 |
| 1866 | Off the southern tip of central Hispaniola (Dominican Republic), Caribbean Sea (ca. 17.50 N, 71.83 W) | 1 | Commercial | Reeves and Mitchell 1988a; Mitchell and Reeves 1988; Bolaños-Jiménez et al. 2014 |
| 1867 | North Atlantic | 1 | Commercial | D. B. Stewart, unpubl. data |
| 1869 | Off Labrador | 1 | Commercial | D. B. Stewart, unpubl. data |
| 1871 | Samana Peninsula, Dominican Republic | 1 | Subsistence | Rodriguez-Demorizi 1960 in Katona et al. 1988 |
| 1874 | North Atlantic | 1 | Commercial | D. B. Stewart, unpubl. data |
| 1884 | Off northwest coast of St. Lucia, West Indies | 1 | Commercial | Reeves 1988; Reeves and Mitchell 1988a; Mitchell and Reeves 1988; Bolaños-Jiménez et al. 2014 |
| 1900 | Trinidad | 1 | Subsistence | di Sciara (in prep.) in Katona et al. 1988 |
| 1902 | Eastport, Maine, USA | 2 | Opportunistic | True 1904; also see Mitchell and Reeves 1988 |
| 1921 | off Florida Keys, Florida, USA | 1 | Opportunistic | MMEP no. STR04968 in Katona et al. 1988 |
| 1924 | Off West Greenland | 1 | Commercial | Smith (Ed.) 2019 |
| 1925 | Trinidad & Tobago | 1 | Subsistence (?) | Reeves and Mitchell 1988a; di Sciara In Litt. in Bolaños-Jiménez et al. 2014 |
| 1934 | Near Hollywood, Florida (ca. 28.03 N) | 1 | Opportunistic | Moore 1953 in Mitchell and Reeves 1988 |
| 1939 | Gulf Stream between Miami Beach and the Bahamas | 1 | Opportunistic | Mowbray 1939 as cited in Moore 1953, p. 139 in Mitchell and Reeves 1988 |
| 1940 | Kap Farvel (Southwest Greenland) | Some | Subsistence | Heide-Jørgensen 1988 |
| 1947 | Southern Trinity Bay, Newfoundland (ca. 47.75N, 53.67 W) | 1 | Commercial | Sergeant and Fisher 1957, Table III; also see Mitchell and Reeves 1988 |
| 1949 | Hyannis, Massachusetts, USA | 1 | Opportunistic | Waters and Rivard 1962 in Katona et al. 1988 |
| 1952 | Upernavik/Aappilattoq, West Greenland | 2 | Subsistence | Heide-Jørgensen 1988 |
| 1954 | Southern Trinity Bay, Newfoundland (ca. 47.75 N, 53.67 W) | 1 | Commercial | Sergeant and Fisher 1957, Table III; also see Mitchell and Reeves 1988; Smith (Ed.) 2019 |

| | | | | |
|---|---|---|---|---|
| 1955 | Grant Suttie Bay, Foxe Basin, Nunavut | 5 (minimum) | Subsistence | Blackadar 1964; Reeves and Mitchell 1988b; Higdon and Ferguson 2014 |
| 1955 | Trinity Bay, Newfoundland | 1 | Commercial | Smith (Ed.) 2019 (but see Mitchell and Reeves 1988) |
| 1957 | Conception Bay, Newfoundland | 3 | Commercial | D.E. Sergeant, in litt. to Economics Br., Dept,. of Fisheries, St. Johns, ABS files in Mitchell and Reeves 1988 |
| 1960 | Kronpronsens Ejland/Disko Bay area, West Greenland | 12 | Subsistence | Heide-Jørgensen 1988 |
| 1961 | Disko Bay area, West Greenland | 2 | Subsistence | Heide-Jørgensen 1988 |
| 1963 | Upernaviup (Disko Bay area), West Greenland | 1 | Subsistence | Heide-Jørgensen 1988 |
| 1964 | Disko Bay, West Greenland | 3 | Subsistence | Heide-Jørgensen 1988 |
| 1964 | South of Newfoundland (ca. 44 N, 55 W) | 1 | Commercial | Mitchell, unpublished data in Mitchell and Reeves 1988; also see Smith (Ed.) 2019 |
| 1964 | Southeast of Nova Scotia (ca. 43.67 N, 59 W) | 1 | Commercial | Mitchell, unpublished data in Mitchell and Reeves 1988; also see Smith (Ed.) 2019 |
| 1965 | Paamiut Isblink (Southwest Greenland) | 5 | Subsistence | Heide-Jørgensen 1988 |
| 1966 | Disko Bay area, West Greenland | 1 | Subsistence | Heide-Jørgensen 1988 |
| 1967 | Southwest Greenland | 1 | Subsistence | Heide-Jørgensen 1988 |
| 1967 | South of Nova Scotia (ca. 44.02 N, 62.43 W) | 1 | Commercial | Mitchell, unpublished data in Mitchell and Reeves 1988; also see Smith (Ed.) 2019 |
| 1968 | Disko Bay area, West Greenland | 2 | Subsistence | Heide-Jørgensen 1988 |
| 1968 | Labrador Sea | 6 | Commercial | Christensen 1982; Oien 1988 |
| 1968 | St. Vincent, Saint Vincent and the Grenadines (Off lee of St. Vincent) | 3 | Subsistence | Caldwell and Caldwell 1975; Katona et al. 1988; Mitchell and Reeves 1988; Bolaños-Jiménez et al. 2014 |
| 1968 | St. Vincent, Saint Vincent and the Grenadines | 3 | Subsistence | Caldwell et al. 1971a; Katona et al. 1988; Mitchell and Reeves 1988; Bolaños-Jiménez et al. |
| 1968 | St. Vincent, Saint Vincent and the Grenadines | 3 | Subsistence | Caldwell et al. 1971b; Caldwell and Caldwell 1975; Katona et al. 1988; Bolaños-Jiménez et al. 2014 |
| 1969 | Disko Bay area, West Greenland | 2 | Subsistence | Heide-Jørgensen 1988 |
| 1969 | Off West Greenland | 1 | Commercial | Smith (Ed.) 2019 |
| 1969 | Labrador Sea | 22 | Commercial | Christensen 1982; Oien 1988 |
| 1969 | Entrance to Clearwater Fiord, Cumberland Sound, Nunavut | 4 | Subsistence | RCMP Game Report, Pangnirtung, 28 August 1970, in Reeves and Mitchell 1988b |
| 1969 | North of Newfoundland (ca. 49.87 N, 52.42 W) | 1 | Opportunistic (?) | Mitchell and Reeves 1988 |
| 1969 | St. Vincent, Saint Vincent and the Grenadines | 4 | Subsistence | Caldwell and Caldwell 1975; Mitchell and Reeves 1988; Bolaños-Jiménez et al. 2014 |

| | | | | |
|---|---|---|---|---|
| 1969 | St. Lucia, West Indies | 1 (+) | Subsistence | Reeves 1988 |
| 1970 | Nuussuaq (Upernavik District)/Disko Bay, West Greenland | 2 | Subsistence | Heide-Jørgensen 1988 |
| 1971 | Paamiut Isblink (Southwest Greenland) | 2 | Subsistence | Heide-Jørgensen 1988 |
| 1971 | St. Vincent, Saint Vincent and the Grenadines | 3 | Subsistence | Caldwell and Caldwell 1975; Katona et al. 1988; Mitchell and Reeves 1988; Bolaños-Jiménez et al. 2014 |
| 1971 | St. Vincent, Saint Vincent and the Grenadines | 9 | Subsistence | Caldwell and Caldwell 1975; Katona et al. 1988; Bolaños-Jiménez et al. 2014 |
| 1971 | Off Bauline, Conception Bay, Newfoundland | 2 | Commercial | Mitchell, unpublished data in Mitchell and Reeves 1988 |
| 1971 | Trinity Bay, Newfoundland | 1 | Commercial | Smith (Ed.) 2019 (but see Mitchell and Reeves 1988) |
| 1971 | Trinity Bay, Newfoundland | 1 | Commercial | Smith (Ed.) 2019 (but see Mitchell and Reeves 1988) |
| 1971 | Off West Greenland | 1 | Commercial | Smith (Ed.) 2019 |
| 1971 | Off West Greenland | 1 | Commercial | Smith (Ed.) 2019 |
| 1972 | Disko Bay area, West Greenland | 2 | Subsistence | Heide-Jørgensen 1988 |
| 1972 | Labrador Sea | 17 | Commercial | Christensen 1982; Oien 1988 |
| 1972 | St. Vincent, Saint Vincent and the Grenadines | 1 | Subsistence | Caldwell and Caldwell 1975; Mitchell and Reeves 1988; Bolaños-Jiménez et al. 2014 |
| 1973 | St. Vincent, Saint Vincent and the Grenadines | 1 | Subsistence | Caldwell and Caldwell 1975; Katona et al. 1988; Mitchell and Reeves 1988; Bolaños-Jiménez et al. 2014 |
| 1974 | St. Vincent, Saint Vincent and the Grenadines | 1 | Subsistence | Caldwell and Caldwell 1975; Katona et al. 1988; Mitchell and Reeves 1988; Bolaños-Jiménez et al. 2014 |
| 1974 | Nuuk/Nanortalik (Southwest Greenland) | 3 | Subsistence | Heide-Jørgensen 1988 |
| 1974 | Off West Greenland | 1 | Commercial | Smith (Ed.) 2019 |
| 1974 | Off West Greenland | 1 | Commercial | Smith (Ed.) 2019 |
| 1975 | Upernavik District, West Greenland | 1 | Subsistence | Heide-Jørgensen 1988 |
| 1977 | Disko Bay area/Kook Isles (Nuuk area), West Greenland | 17 | Subsistence | Heide-Jørgensen 1988 |
| 1977 | Kekertelung Island, Cumberland Sound, Nunavut | 14 | Subsistence | Mitchell 1979; Davis et al. 1980; Mitchell and Reeves 1988; Reeves and Mitchell 1988b |
| 1978 | Baker Lake, Nunavut | 1 | Subsistence | Davis et al. 1980; Kayuryuk and Innakatsik 1982; Sergeant 1986; Reeves and Mitchell 1988b |
| 1980 | Nuuk, West Greenland | 2 | Subsistence | Heide-Jørgensen 1988 |
| 1981 | Offshore Davis Strait | 1 | Subsistence | Heide-Jørgensen 1988 |
| 1982 | Qassimiut (Southwest Greenland) | 1 | Subsistence | Heide-Jørgensen 1988 |

| | | | | |
|---|---|---|---|---|
| 1982 | St. Vincent, Saint Vincent and the Grenadines | 1 | Subsistence | Price 1982 in Katona et al. 1988; Price 1983 in Bolaños-Jiménez et al. 2014 |
| 1982 | St. Vincent, Saint Vincent and the Grenadines | 2 | Subsistence | Price 1982 in Katona et al. 1988; Price 1983 in Bolaños-Jiménez et al. 2014 |
| 1984 | Disko Bay area, West Greenland | 1 | Subsistence | Heide-Jørgensen 1988 |
| Pre-1985 | St. Vincent, Saint Vincent and the Grenadines | 6 | Subsistence | Price 1985 |
| 1986 | Sisimiut and Maniitsoq Districts, West Greenland | 17 | Subsistence | Heide-Jørgensen 1988 |
| 1986 | Columbia | 1 | Retaliatory | Álvarez-León 2002 in Bolaños-Jiménez et al. 2014 |
| 1987 | Nuuk, West Greenland | 1 | Subsistence | Heide-Jørgensen 1988 |
| 1987 | Trinidad & Tobago | 1 | Fishing gear entanglement | Ottley et al. 1988 in Bolaños-Jiménez et al. 2014; Vidal et al. 1994 |
| 1994 | St. Vincent, Saint Vincent and the Grenadines | 2 | Subsistence | Bolaños-Jiménez et al. 2014 |
| 1996 | West Greenland | 3 | Subsistence | NAMMCO 2019 |
| 1996 | St. Vincent, Saint Vincent and the Grenadines | 5 | Subsistence | Sutty, unpublished data in Bolaños-Jiménez et al. 2014 |
| 1997 | West Greenland | 4 | Subsistence | NAMMCO 2019 |
| 1998 | West Greenland | 1 | Subsistence | NAMMCO 2019 |
| 1999 | West Greenland | 6 | Subsistence | NAMMCO 2019 |
| 2000 | West Greenland | 1 | Subsistence | NAMMCO 2019 |
| 2000 | St. Vincent, Saint Vincent and the Grenadines | 3 | Subsistence | Sutty, unpublished data in Bolaños-Jiménez et al. 2014 |
| 2001 | West Greenland | 2 | Subsistence | NAMMCO 2019 |
| 2002 | West Greenland | 21 | Subsistence | NAMMCO 2019 |
| 2003 | West Greenland | 3 | Subsistence | NAMMCO 2019 |
| 2004 | West Greenland | 12 | Subsistence | NAMMCO 2019 |
| 2005 | West Greenland | 2 | Subsistence | NAMMCO 2019 |
| 2007 | West Greenland | 3 | Subsistence | NAMMCO 2019; also see Altherr and Hodgins 2018 |
| 2007 | St. Vincent, Saint Vincent and the Grenadines | 1 | Subsistence | Fielding 2010 and pers. comm. in Bolaños-Jiménez et al. 2014 |
| 2008 | West Greenland | 26 | Subsistence | NAMMCO 2019; also see Altherr and Hodgins 2018 |
| 2008 | St. Vincent, Saint Vincent and the Grenadines | 2 | Subsistence | Fielding 2010 and pers. comm. in Bolaños-Jiménez et al. 2014 |
| 2008 | St. Vincent, Saint Vincent and the Grenadines | 3 | Subsistence | Fielding 2010 and pers. comm. in Bolaños-Jiménez et al. 2014 |
| 2008 | St. Vincent, Saint Vincent and the Grenadines | 1 | Subsistence | Fielding 2010 and pers. comm. in Bolaños-Jiménez et al. 2014 |
| 2008 | St. Vincent, Saint Vincent and the Grenadines | 3 | Subsistence | Fielding 2010 and pers. comm. in Bolaños-Jiménez et al. 2014 |
| 2008 | St. Vincent, Saint Vincent and the Grenadines | 2 | Subsistence | Fielding 2010 and pers. comm. in Bolaños-Jiménez et al. 2014 |
| 2008 | St. Vincent, Saint Vincent and the Grenadines | 1 | Subsistence | Fielding 2010 and pers. comm. in Bolaños-Jiménez et al. 2014 |
| 2009 | West Greenland | 9 | Subsistence | NAMMCO 2019; also see Altherr and Hodgins 2018 |
| 2010 | West Greenland | 14 | Subsistence | NAMMCO 2019; also see Altherr and Hodgins 2018 |

| | | | | |
|---|---|---|---|---|
| 2011 | West Greenland | 34 | Subsistence | NAMMCO 2019; also see Altherr and Hodgins 2018 |
| 2011 | St. Vincent, Saint Vincent and the Grenadines | 1 | Subsistence | Bolaños-Jiménez et al. 2014 |
| 2012 | West Greenland | 32 | Subsistence | NAMMCO 2019; also see Altherr and Hodgins 2018 |
| 2013 | West Greenland | 27 | Subsistence | NAMMCO 2019; also see Altherr and Hodgins 2018 |
| 2014 | West Greenland | 10 | Subsistence | NAMMCO 2019; also see Altherr and Hodgins 2018 |
| 2015 | West Greenland | 16 | Subsistence | NAMMCO 2019; also see Altherr and Hodgins 2018 |
| 2015 | St. Vincent, Saint Vincent and the Grenadines | 4 | Subsistence | Gaworecki 2015; Altherr and Hodgins 2018 |
| 2016 | West Greenland | 12 | Subsistence | NAMMCO 2019; also see Altherr and Hodgins 2018 |
| 2017 | West Greenland | 6 | Subsistence | NAMMCO 2019 |
| 2017 | St. Vincent, Saint Vincent and the Grenadines | 2 | Subsistence | Gibbens 2017; Altherr and Hodgins 2018 |
| 2018 | St. Vincent, Saint Vincent and the Grenadines | 3 | Subsistence | Chance 2018; Altherr and Hodgins 2018 |
| 2019 | Near Beaumont, Long Island, northeast Newfoundland | 1 | Fishing gear entanglement | CBC 2019 |
